# Recombination, truncation and horizontal transfer shape the diversity of cytoplasmic incompatibility patterns

**DOI:** 10.1101/2025.01.06.631454

**Authors:** Alice Namias, Julien Martinez, Iliana Boussou, Kevin Terretaz, Will Conner, Fabienne Justy, Patrick Makoundou, Marco Perriat-Sanguinet, Pierrick Labbé, Mathieu Sicard, Frederic Landmann, Mylène Weill

## Abstract

*Wolbachia* are endosymbiotic bacteria inducing various reproductive manipulations of which cytoplasmic incompatibility (CI) is the most common. CI leads to reduced embryo viability in crosses between males carrying *Wolbachia* and uninfected females or those carrying an incompatible symbiont strain. In the mosquito *Culex pipiens*, the *Wolbachia w*Pip causes highly complex crossing patterns. This complexity is linked to the amplification and diversification of the CI causal genes, *cidA* and *cidB*, with polymorphism located in the CidA-CidB interaction regions. We previously showed correlations between the identity of gene variants and CI patterns. However, these correlations were limited to specific crosses, and it is still unknown whether *c*id gene polymorphism in males’ and females’ *Wolbachia* can explain and predict the wide range of crossing types observed in *C. pipiens*. Taking advantage of a new method enabling full-gene acquisition, we sequenced complete *cid* repertoires from 45 *w*Pip strains collected worldwide. We demonstrated that the extensive diversity of *cid* genes arises from recombination and horizontal transfers. We uncovered further *cidB* polymorphism outside the interface regions and strongly correlated with CI patterns. Most importantly, we showed that in every *w*Pip genome, all but one *cidB* variant are truncated. Truncated *cidB*s located in palindromes are partially or completely deprived of their deubiquitinase domain, crucial for CI. The identity of the sole full-length *cidB* variant seems to dictate CI patterns, irrespective of the truncated *cidBs* present. Truncated CidBs exhibit reduced toxicity and stability in *Drosophila* cells, which potentially hinders their loading into sperm, essential for CI induction.

## Introduction

*Wolbachia* are endosymbiotic alpha-proteobacteria that infect nematode and arthropod species. In the latter, they are most often reproductive parasites, causing various reproductive manipulations of which cytoplasmic incompatibility (CI) is the most common. In its simplest form, CI is defined by elevated embryo mortality in crosses between *Wolbachia*-infected males and uninfected females. CI can also occur in crosses between individuals infected by so-called incompatible *Wolbachia*. CI is formalized as a modification-rescue (or *mod-resc*) system (Werren 1997), where the modification factor affects paternal DNA, causing developmental defects and leading to embryo death, unless the appropriate rescue factor is present in the egg (Werren et al. 2008). Although CI has been studied for 70 years (Laven 1951; Laven 1967), the key CI genes, named *cif*, were discovered in 2013 (Beckmann and Fallon 2013), and their role in CI was only demonstrated in 2017 (Beckmann et al. 2017; LePage et al. 2017). Ten phylogenetic groups (or types) of *cif* genes have been described in all *Wolbachia* strains depending on the functional domains on the *cifB* gene (LePage et al. 2017; Lindsey et al. 2018; Bing et al. 2020; Tan et al. 2024) and named type I-X. In the *Wolbachia w*Pip infecting *Culex pipiens* mosquitoes, genes essential for determining CI patterns are from type I and named *cidA* and *cidB*. *cid* stands for CI deubiquitinase (DUB), a DUB domain present in the downstream part of the *cidB* gene. This DUB domain has been shown to be key for CI (Beckmann et al. 2017). First thought to be responsible for the toxicity at the heart of CI, recent studies have shown that it is rather involved in the loading of the CidB proteins into the sperm – a key step for CI induction (Horard et al. 2022; Terretaz et al. 2023). One explanatory model for CI particularly suited to *Culex pipiens* is the toxin- antidote model involving *cidA* as the antidote and *cidB* as the toxin. Compatibility results from binding between the toxin and the antidote, while incompatibility occurs when no antidote binds (and thus neutralizes) the toxin (Namias et al. 2022).

*Wolbachia*-induced CI has been extensively studied in *Culex pipiens*, revealing highly complex CI patterns (Laven 1967; Atyame et al. 2014; Namias et al. 2022). These patterns were long puzzling, as no polymorphism could be found in *w*Pip genomes using standard multilocus sequence typing (MLST) *Wolbachia* markers (Baldo et al. 2006). An important step in the understanding of CI patterns was the discovery of five phylogenetic groups within *w*Pip, named *w*Pip-I to *w*Pip-V, enabled by the use of hypervariable *w*Pip-specific markers (Atyame, Delsuc, et al. 2011). Briefly, crosses involving males and females infected with *w*Pip strains belonging to the same *w*Pip group were largely compatible, while crosses involving different *w*Pip groups had unpredictable outcomes (Atyame et al. 2014). This high CI complexity led both experimental and modeling studies to conclude about the presence of several toxins and antidotes in each *Wolbachia* genome (Atyame, Duron, et al. 2011; Nor et al. 2013). Indeed, it was later shown that *cid* genes were amplified and diversified in *w*Pip genomes (Bonneau, Atyame, et al. 2018). For both *cidA* and *cidB*, polymorphism was found to be restricted to two specific regions, named upstream and downstream (Bonneau, Atyame, et al. 2018). In the previously investigated *w*Pip (Bonneau, Atyame, et al. 2018; Bonneau et al. 2019; Sicard et al. 2021), up to six different *cidA* copies and four different *cidB* copies were reported within a single genome. The different sequences of the *cid* genes were named variants, and the set of variants present within a given genome was called a repertoire. To date, around 30 different *cidA* variants and 20 different *cidB* variants have been described (Bonneau, Atyame, et al. 2018; Bonneau, Landmann, et al. 2018; Bonneau et al. 2019; Namias et al. 2023; Namias et al. 2024). Variable upstream and downstream regions were predicted (Bonneau, Atyame, et al. 2018) and then confirmed by crystal structure (Xiao et al. 2021; Wang et al. 2022) to be involved in the binding interface between CidA and CidB proteins. Focusing on crosses between *w*Pip-IV males from 245 different *Culex pipiens* lines and *w*Pip-I females, we revealed a strong correlation between the presence or absence of a specific recombinant *cidB* variant (*cidB*-IV-2) and CI crossing phenotype variations (compatible/incompatible), although some rare discrepancies remained unexplained (Bonneau, Atyame, et al. 2018; Bonneau et al. 2019). In a recent study of CI microevolution in a laboratory isofemale line, we found that the rapid loss of a *cidA* variant in females’ repertoire perfectly matched the observed shift in compatibility patterns. Females with *Wolbachia* without this antidote, which is recombinant in the binding interface region, were unable to rescue CI induced by *w*Pip-IV males with the recombinant variant *cidB*-IV-2 (Namias et al. 2024). These results demonstrate the major role played by CidA-CidB binding interface polymorphism in CI pattern diversity, thus strongly supporting the fact that binding between CidA and CidB is central in CI, as affirmed by the toxin-antidote model (Namias et al. 2022).

Until the present study, repertoires were obtained by sequencing PCR products, targeting only the variable regions of the *cid* genes (Bonneau, Atyame, et al. 2018) involved in the CidA/CidB binding interface. This was performed using cloning and Sanger sequencing (Bonneau, Atyame, et al. 2018; Bonneau, Landmann, et al. 2018; Bonneau et al. 2019; Sicard et al. 2021), and more recently, by direct Nanopore sequencing (Namias et al. 2023; Namias et al. 2024). Targeted acquisition is easier than full-genome acquisition and genome assembly, which are troublesome in *w*Pip. Indeed, *cid* genes are located in highly repeated prophagic regions, resulting in strong discordances between the two *w*Pip reference draft genomes in terms of *cid* repertoires. The *w*Pip-Pel genome (Klasson et al. 2008) is not fully circular and contains a single pair of *cid* genes, an observation that is unlikely given the high number of *cid* gene copies observed in all *w*Pip studied to date (Bonneau, Atyame, et al. 2018; Bonneau, Landmann, et al. 2018; Bonneau et al. 2019; Sicard et al. 2021; Namias et al. 2023; Namias et al. 2024). Nevertheless, the *w*Pip-JHB genome, likewise fragmented (Salzberg et al. 2009), contains several *cid* pairs (putatively three (Lindsey et al. 2018; Martinez et al. 2021)), but exhibits several loss-of-function mutations as well as truncations of the *cidB* 3’ end (Martinez et al. 2021). Open reading frame-disrupting mutations of *cif* genes are common across *Wolbachia* genomes, particularly in the case of *cifB* (Martinez et al. 2021), although their effect on protein function has not yet been investigated in detail.

To investigate if such truncated *cidB*s were present in other *w*Pip genomes, we Nanopore-sequenced two genomes. Both exhibited truncated *cidB* variants and lacked their DUB domains. These new results prompted us to develop a new sequencing strategy that gave access to all *cidA* and *cidB* polymorphism located both inside and outside the binding interface regions. In the present study, we acquired the full *cid* repertoires of *w*Pip from 45 phylogenetically and geographically diverse *C. pipiens* lines (41 isofemale lines and 4 individuals from natural populations). We showed that almost all polymorphism in the CidA/CidB interface region resulted from recombinations of numerous sequence blocks rather than mutations. We also demonstrated that in each repertoire, only a single full-length *cidB* variant was found, as all the other variants were truncated. The identity of the sole full-length *cidB* variant seems to dictate the CI patterns, irrespective of the truncated *cidB*s present in the same genome. Compared to full-length CidB proteins, their truncated counterparts exhibit reduced toxicity and stability when expressed in Drosophila cells, thus potentially hindering the loading in sperm required to contribute to the CI mechanism.

## Results

### High polymorphism in CidA/CidB interface regions results from recombination

We acquired the repertoire of *cidA* and *cidB* genes (corresponding to the interface regions) for 73 individuals from 41 different isofemale lines and 4 field populations (Table S1). Whenever possible, two mosquitoes were analyzed per line, giving mostly identical variants (Table S2 and S3). We named the new variants after the nomenclature updated in (Namias et al. 2024). Including the previously sequenced variants, we now had a total of 56 distinct *cidA* variants (Table S2) and 55 distinct *cidB* variants (Table S3). Using recombination analysis methods (RDP4, (Martin et al. 2015)), we showed that most of the polymorphism resulted from recombination: we identified 15 recombination blocks (i.e., sets of adjacent single nucleotide polymorphisms (SNPs) that are always inherited together) in *cidA* and 23 in *cidB*, which were validated with at least four distinct recombination analysis methods. While a high number of recombination blocks were found, very few standalone SNPs were not included in a recombination block (5 for *cidA* and 2 for *cidB*) (Fig 1 and Fig S1).

**Figure 1.**
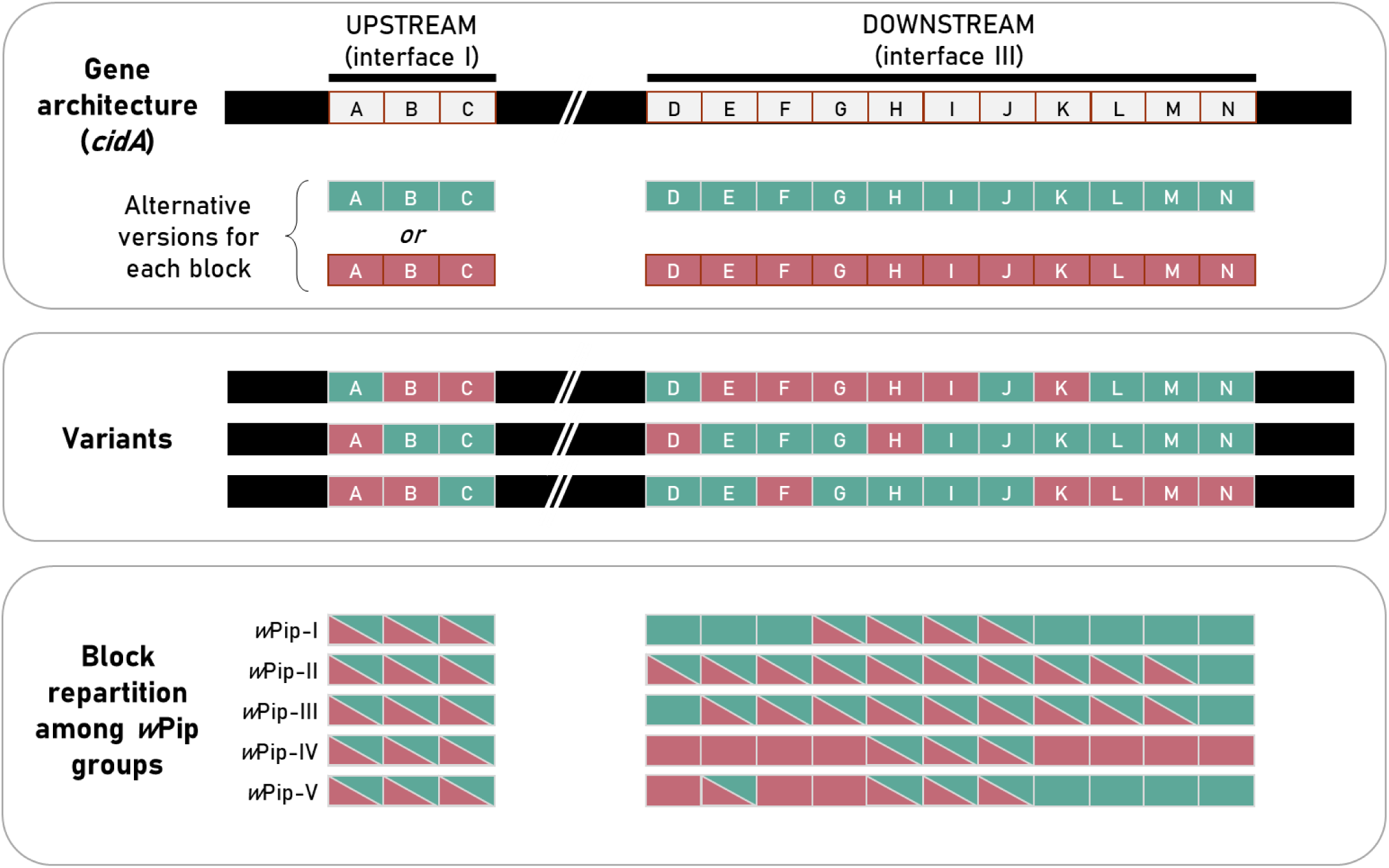
Recombinant *cidA* variants with common architecture. Monomorphic gene regions are shown in black. Polymorphic regions are composed of several recombination blocks, named A to 0. These blocks are always in the same order, with each having two distinct alternative versions, shown here in blue and red. A variant is the assembly of different versions of each block. The lowest panel shows the repartition of block versions among *w*Pip groups. If the two versions of a block are found in a given group, the block contains triangles of both colors, whereas if a single block version is found, the cell shows a single color. Colors are arbitrarily determined, and versions of a given color from different blocks are not more likely to be found together.

CidA and CidB proteins interact head to tail through three interaction interfaces (Xiao et al. 2021; Wang et al. 2022). Interaction interface I, which corresponds to the upstream region of *cidA* and the downstream region of *cidB,* shows little polymorphism, with four possible regions for *cidA* (alpha, beta, gamma, and delta) and three for *cidB* (1, 2, and 3). By contrast, interaction interface III, which corresponds to the downstream region of *cidA* and the upstream region of *cidB,* is highly polymorphic, with 23 and 36 distinct regions for *cidA* and *cidB*, respectively (Table S2 and S3, Fig 1 and Fig S1). Finally, interaction interface II is not polymorphic.

For both *cidA* and *cidB*, all recombination blocks always show exactly two alleles, meaning that no position has three or more possible nucleotides in the full set of sequences. All observed *cid* variants are made by combining one of the two different alleles of each block (Fig 1 and Fig S1 for *cidA* and *cidB*, respectively). Although *w*Pip phylogenetic groups are key to CI patterns (Atyame et al. 2014), all block alleles are shared among groups, apart from a few exceptions specific to *w*Pip-IV (blue allele of blocks P and Q for *cidB*, and red allele of block O for *cidA*; Fig 1 and Fig S1). Phylogenetic groups *w*Pip- II and *w*Pip-III show the highest sequence diversity for both *cidA* and *cidB*, as almost all possible block alleles are found in these groups (Fig 1 and Fig S1). Whereas block alleles are largely shared among groups, variants (i.e., combinations of blocks) are *w*Pip-group specific. This is especially clear for *cidB*, as none of the 55 *cidB* variants were shared between *w*Pip groups. For *cidA*, only 6 out of 56 variants were shared between *w*Pip groups: two were shared exclusively by *w*Pip groups II and III, three exclusively by *w*Pip groups II and V, and one by *w*Pip groups II, III, and V.

### A unique full-length CidB protein is key to CI crossing types

#### Palindromic *cidA-cidB* tandems contain truncated *cidB* variants

One of the *w*Pip draft reference genomes, *w*Pip-JHB, shows two putative pairs of *cidA-B* genes (scaffold Genbank accession: DS996944.1) as well as a partially sequenced cidB 3’ end (scaffold Genbank accession: DS996943.1). The *cidBs* in *cidA-B* pairs recovered from this genome assembly are truncated, lacking the 3’ end of the gene (no ambiguous bases at the truncation sites, suggesting that these truncations are real) (Salzberg et al. 2009; Martinez et al. 2021). To investigate these truncations, we acquired Nanopore long reads corrected by Illumina sequencing for two *w*Pip strains: *w*Pip-Tunis (*w*Pip-I) and *w*Pip-Harash (*w*Pip-IV). Nonetheless, we were unable to assemble the genome due to the very high presence of repeated elements, particularly around *cid* genes. However, we analyzed the long reads containing *cid* genes, finding that *cidA* and *cidB* genes were always in tandem, with *cidA* upstream of *cidB*. Tandems were found alone (similar to what is observed in the *w*Pip-Pel genome) or organized by pairs in palindromes (*cidA-cidB-cidB-cidA*). All *cidB* located in these palindromes were truncated, lacking their 3’ end, whereas all the *cidA* variants were complete.

Two distinct palindromes were observed: the first (P1) was common to *w*Pip-Tunis and *w*Pip-Harash, but the second (P2) was only found in *w*Pip-Harash (Fig 2). P1 and P2 had a common truncated *cidB*, truncated at 2,466 base pairs (bp) corresponding to an 822 amino-acid protein, referred to hereafter as *cidB-*TrA. In P1, the other truncated *cidB* (TrB) was composed of (i) the first 2,381 bp of *cidB*, (ii) an in-frame “insert” of 551 bp, and (iii) the last 83 bp corresponding to the reverse-complement of the overlapping *cidB*-TrA 3’ end, resulting in an open reading frame of 3,015 bp (1,005 amino acids). In P2, the other truncated *cidB* (*cidB*-TrC) was comprised of (i) the first 2,958 bp of *cidB*, (ii) an in-frame “insert” of 25 bp, and (iii) the same last 83 bp of *cidB*-TrA, resulting in an open reading frame of 3,066 bp (1,022 amino acids). The so-called inserts were defined by comparison with the reference genome *w*Pip-Pel (Klasson et al. 2008). The three distinct truncated CidB proteins lacked all or part of their DUB domain (Fig 2). Truncated *cidB* variants identified in the *w*Pip-JHB genome (Martinez et al. 2021) were organized as a P1 palindrome. Since P1 was found in three distinct genomes (Tunis, Harash, and JHB), we designed a P1-specific PCR (Table S4) and tested for the presence of truncated *cid* variants in nine laboratory strains from diverse *w*Pip groups and geographic origins. We identified the P1 palindrome in all the tested strains (Table S1).

**Figure 2.**
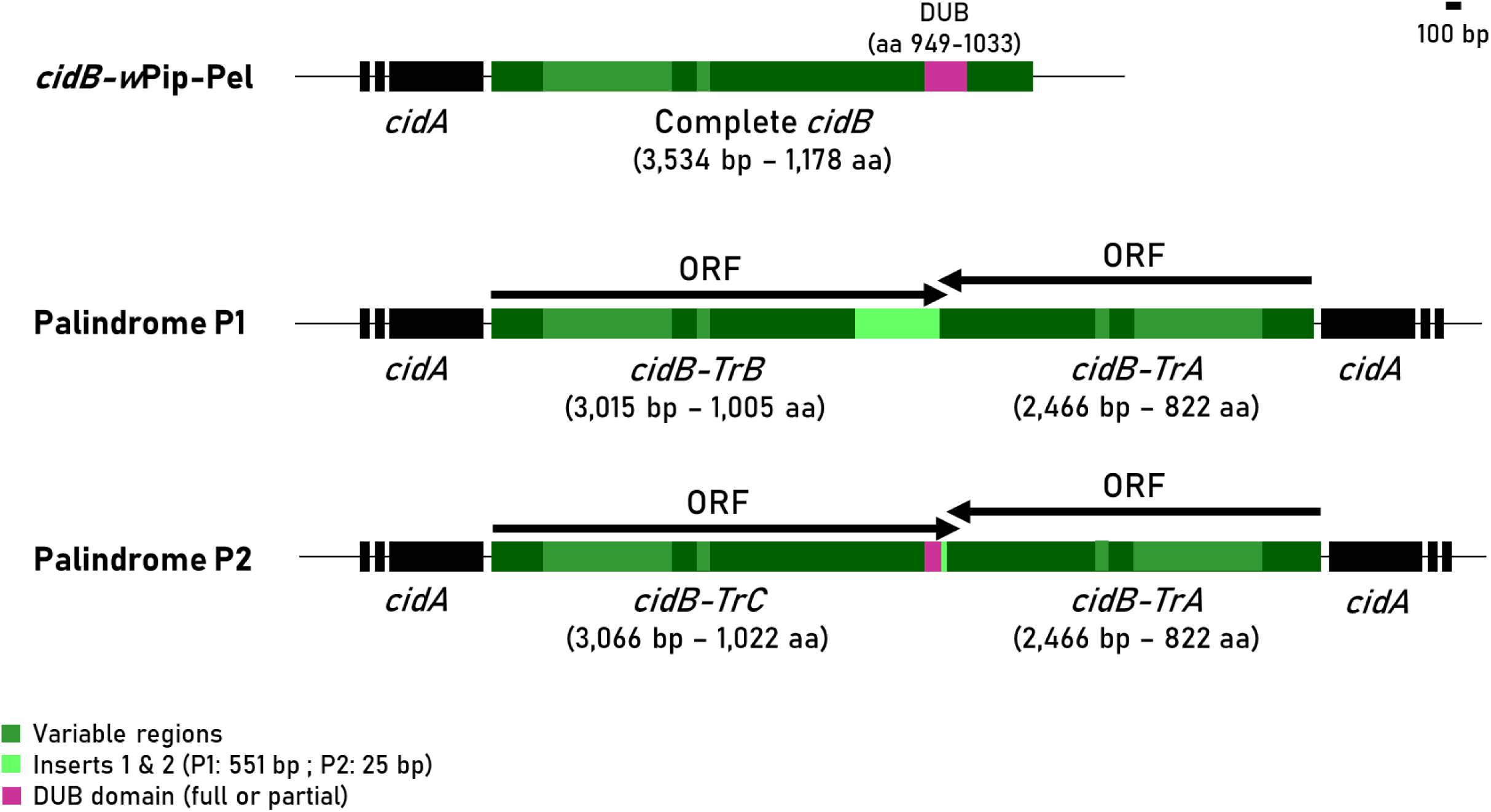
Palindromes observed in the genome long reads of the strains Tunis (*w*Pip-I) and Harash (*w*Pip-IV). The complete *cidB* described in the *w*Pip-Pel genome is given here for comparative purposes. Two distinct palindromes P1 and P2 were found with a *cidA-cidB-cidB-cidA* structure, where the first two genes are sense-orientated and the following two are in opposite directions. For both palindromes, the two *cidB* are truncated. They share a common *cidB*, *cidB-*TrA, truncated at 2,466 base pairs (bp). *cidB-TrB* (found in P1) and *cidB*-*TrC* (found in P2) are chimeric open reading frames (ORFs) that contain part of the “classic” *cidB* (2,381 bp and 2,958 bp for *cidB-TrB* and *cidB*-*TrC*, respectively), an insert (551 bp and 25 bp, respectively) and part of the *cidB*-*TrA* (83 bp for both). Overall, the *cidB-TrB* ORF is 3,015 bp, while that of *cidB-TrC* is 3,066 bp.

To investigate the evolutionary origin of *w*Pip palindromes, we screened publicly available *Wolbachia* genomes and identified a total of 468 *cifA-B* gene pairs, including 121 type I *cid* homologs. The majority contained distinct *cif* gene types and often several copies of a given type regardless of host taxonomy (Table S5). Of 138 screened *Wolbachia* strains, only the *Wolbachia* found in the moth *Rhopobota naevana* (tentatively named *w*Naev, (Vancaester and Blaxter 2023)) exhibited a *cif* palindrome (Table S5, Fig S2A). Species closely related to *w*Pip did not display similar palindromic structures, suggesting that this structure is not ancestral to *w*Pip-like strains. Therefore, it could have emerged within the *w*Pip genome or been acquired through horizontal transfer from a distantly related strain. The palindrome observed in *w*Naev involved distantly related *cifs*, a type I divergent from *w*Pip *cids* and a type V (Fig S2B), while the palindromes from *w*Pip involves two *cids* (type I). These palindromes could result from distinct events, or one of them could have resulted from a rearrangement of the other palindrome. To address this question, we first tried to understand how P1 was formed, since it is common to all the tested *w*Pip strains. Blasting the *cidB-TrB* gives a perfect hit with the first 3 kb of a non-palindromic *cidB* gene present in the genome of *w*Naev (*cidB-Naev*, Fig 3; pair 4, Fig S2). Interestingly, the 3’ sequence of this *cidB* was also found in *w*Pip-Pel genome at positions 1,372,335 to 1,372,841 (WP_1292) (Fig 3). The complete sequence of the *cidB-Naev* pair 4 (type I) was thus found in *w*Pip genomes but split into two parts. This strongly suggests a horizontal transfer of *cidB-Naev* (or a similar strain not yet sequenced) to an ancestral *w*Pip before *w*Pip diversification into *w*Pip groups. This *cidB* could have been directly rearranged during its acquisition by *w*Pip, resulting in the observed palindrome, or been transferred and later rearranged (Fig 3).

**Figure 3.**
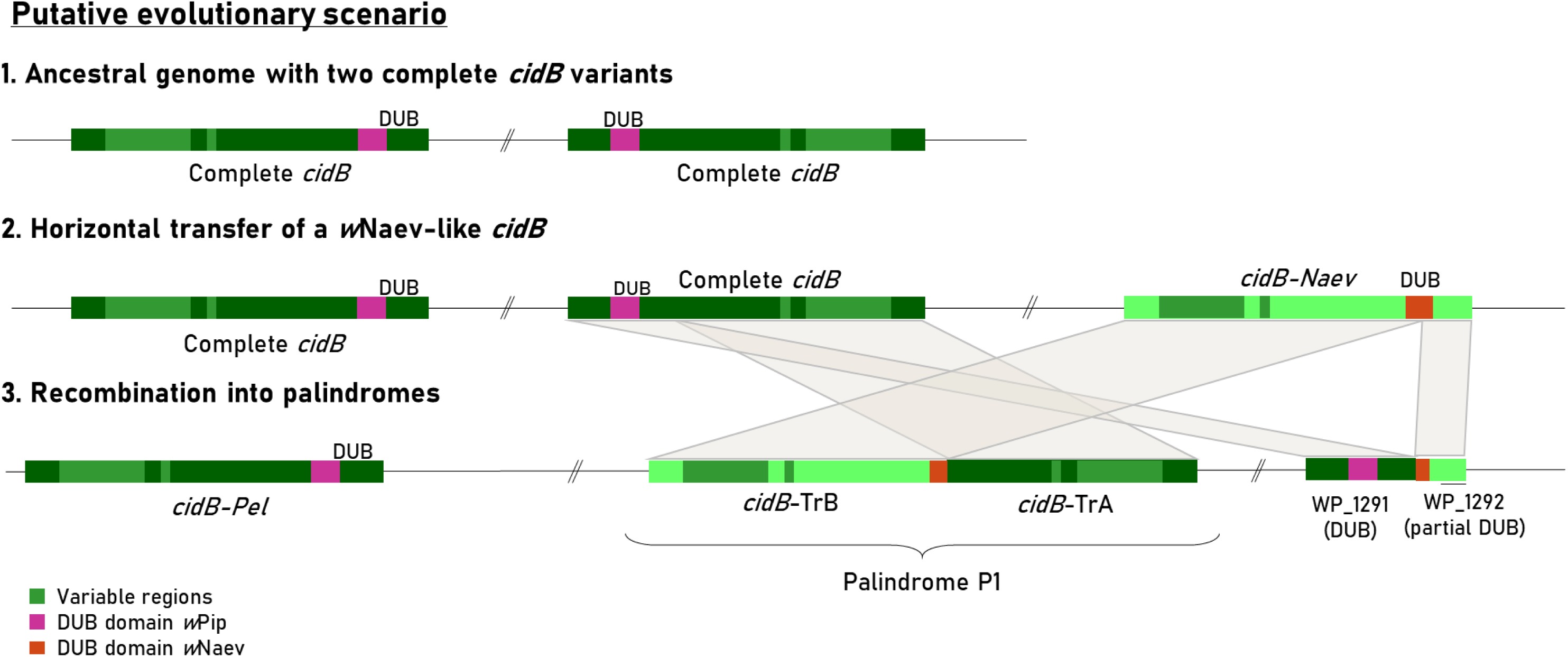
Evolutionary scenario for the evolution of palindromes in *w*Pip. A three-step evolutionary scenario could explain the current genomic architecture of the *cid* genes for the *w*Pip and *w*Naev genomes: all *w*Pip repertoires acquired so far include at least a complete *cidB* along with a palindromic structure containing two truncated *cidB*. All sequenced genomes also have two further *cidB* modules, the WP_1291 and WP_1292 open reading frames (ORFs). In *w*Naev genomes, 10 *cidB* were recovered (Fig S2), one of which belongs to pair 4 and is represented here in light green. *w*Pip *cid* architecture could result from an ancestral genome with two complete *cidB* copies, which underwent a horizontal transfer of a *cidB* from *w*Naev or a genome with a similar *cidB*, followed by a recombination into a palindromic structure.

For palindrome P2, the insert was smaller (25 pb) but part of insert 1 and thus also displayed *cidB-Naev* (among others) as a best hit. This, along with the fact that both palindromes share *cidB-TrA*, this could suggest that P2, which is not fixed in *w*Pip, derives from a rearrangement of P1 in some *w*Pip strains as opposed to an independent event. This rearrangement would have replaced *cidB-TrB* by *cidB*-*TrC*, thus keeping a small piece of the insert.

#### All but one *cidB* variant are truncated

Our previous strategy to characterize *cid* repertoires focused only on the binding interface regions, with PCR products encompassing both *cid* upstream and downstream regions. This PCR amplification of polymorphic regions thus targeted both truncated and complete *cidB* variants but did not discriminate them. We thus designed new generic PCR primers to amplify *cidA* and *cidB* full-length genes (Table S4, Fig S3). To sequence these PCR products, we used a new rapid and efficient repertoire acquisition method based on Nanopore sequencing (Namias et al. 2023). We subsequently label as “short” repertoires the *cid* repertoires obtained by amplifying solely the “polymorphic regions” and as “long” *cid* repertoires those obtained by amplifying the full-length *cid* sequence (Fig S3).

Short and long *cidA* were acquired for approximately 20 lines, and no discrepancies were found between short and long repertoires, further supporting the lack of truncations in the *cidA* genes. We then acquired only short *cidA* repertoires, hereafter called “*cidA* repertoires.” Two to five different *cidA* variants were found depending on the *w*Pip strains (Table S2).

We acquired the short and long *cidB* repertoires for the 73 individuals (Table S3). A comparison of the short and long *cidB* repertoires revealed their considerable difference: while short *cidB* repertoires exhibited one to five different *cidB* variants, the long repertoires were always composed of a unique full-length variant in all 73 sequenced repertoires (Table S3) regardless of the *w*Pip groups. We performed direct Sanger sequencing of the long *cidB* PCR products on 12 strains to verify this result (Table S1). Inspection of the electropherograms showed no multi-peaks, indicating a lack of within-individual polymorphism. This further corroborates the existence of a single full-length *cidB* variant within the investigated *w*Pip genomes (all other *cidB* being truncated). However, this result does not rule out the presence of several identical full-length copies.

#### Toxicity of the truncated *cidB* variants

The *cidB-*TrA and *cidB-*TrB truncations remove partially (TrB) or totally (TrA) the DUB domain of the CidB. This domain was previously shown to be crucial for CidB stability *in vivo* and *in cellulo* (Horard et al. 2022). Despite its reduced stability, a DUB deletion mutant of CidB-IV-a-2 is still able to induce apoptosis when expressed in *Drosophila* S2R+ cells (Terretaz et al. 2023).

To study the impact of these truncations on CidB activity, we generated fluorescent fCidB-IV-a-2-TrA and fCidB-IV-a-2-TrB thanks to a fusion to the superfolder GFP (sfGFP), co-expressed with a fluorescent mKate2 transfection reporter through a T2A self-cleaving peptide (Fig S4A, (Terretaz et al. 2023)).

We first tested the *in cellulo* toxicity of these constructs compared to the full-length fCidB-IV-a-2 variant (full-length CidB, hereafter named fCidB). To this end, we performed time-lapse microscopy of the transfected *Drosophila* S2R+ cells to establish their individual cellular fate (apoptosis, mitosis, or no specific event — interphase — over 48h) (Fig 4A and Table S6A). As previously established, expression of fCidB leads to cell death and prevents mitosis (only one mitosis observed among 1005 observations, *f_m_*= 0.001 [0.001-0.005], with *f_m_* being the frequency of mitosis events observed over the sum of mitosis and apoptosis, and its 95% confidence interval in brackets). This phenotype is rescued by the co-expression of the antidote fCidA-IV-delta(1)-1 (hereafter named fCidA), which restores mitosis (*f_m_*= 0.80 [0.76-0.83]). While cell division is almost never observed with the full-length CidB that blocks DNA replication during the S-phase (Horard et al. 2022; Terretaz et al. 2023), truncated CidB allows mitosis to occur (*f_m_*= 0.2 [0.17-0.23] and *f_m_*= 0.27 [0.23-0.32] for CidB-TrA and CidB-TrB, respectively; Fig 4A). Mitosis frequency was similar for fCidB-TrA and fCidB-TrB (GLMM to consider the replicate effect, LRT, *χ^2^* = 1.75, *p* = 0.19) but significantly lower than fCidA-fCidB (rescue) and significantly higher than fCidB (GLMM, LRT, *χ^2^* = 1192, *p* < 0.001). Although toxic, truncated CidB is less so than the full-length CidB *in cellulo*.

**Figure 4.**
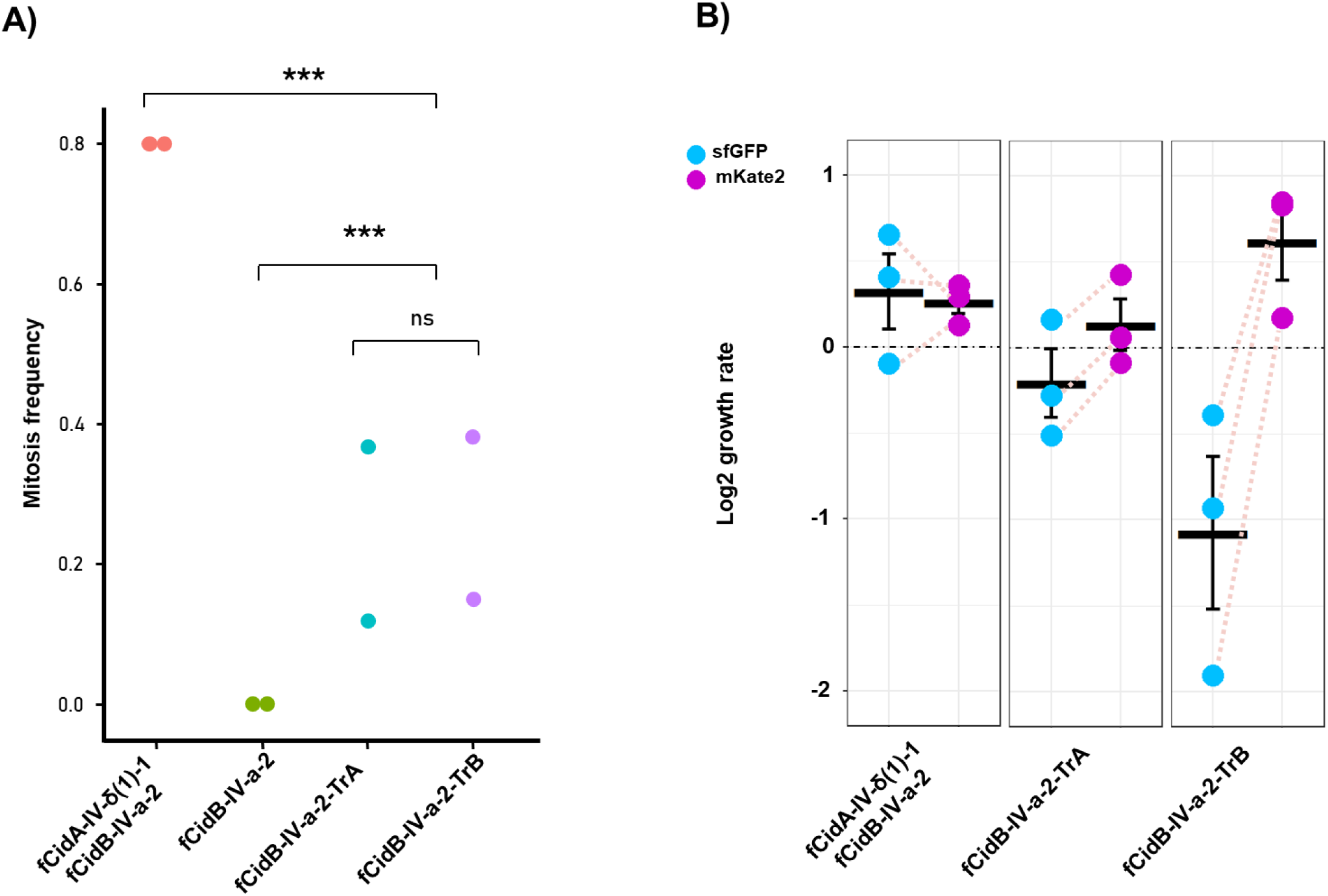
Truncated CidB variants less toxic and less stable than the full-length CidB toxin. (A) Mitosis frequency over mitosis and apoptosis observed for each construct after the transfection of S2R+ cells over 48h of time-lapse microscopy. Each transfection experiment was performed twice independently, and the number of transfected cells observed is >1000. (B) Flow cytometry analyses of S2R+ cell growth from day 2 to day 4 after transfection in three independent transfection experiments. The blue and pink dots represent log2(sfGTP) and log2(mKate2) for each construct, while the associated measures are linked for each biological replicate.

Additional confocal microscopy observations of transfected S2R+ cells indicate that both truncated variants localize to the nucleus, similar to the complete variant fCidB (Fig S4B). Both truncated variants either lead to apoptosis or allow mitosis, decorating the condensed chromosomes in the latter case (Fig S4B). In addition, fCidB-TrA and fCidB-TrB variants interact with fCidA in co-expression experiments. In this case, when the toxin is neutralized by its CidA cognate partner, both effectors remain in the cytoplasm in interphase and then go on to decorate the chromatin during mitosis (Fig S4C). Hence, these truncations do not seem to affect the rescuing process and thus the toxin-antidote interaction.

To evaluate the impact of these truncations on CidB stability over time, we followed the sfGFP fluorescence associated with the variants in cases in which mitoses were observed (i.e., all except the fully toxic fCidB, as evaluating stability requires cell survival). We also took advantage of the free mKate2 fluorescence used as a transfection marker to perform flow cytometry analyses. We monitored fluorescence at days 2 and 4 post-transfection, expressed as the log_2_(sfGFP)/log_2_(mKate2) ratio in a cell growth assay (Terretaz et al. 2023). We adopted the following rationale: if a truncated CidB variant is stable, independently of its toxicity, its associated sfGFP fluorescence must be equivalent to that of the co-expressed free mkate2; however, if it is unstable, its associated GFP fluorescence disappears from the transfected cells, while the free mKate2 fluorescence remains present. We first found that the log_2_(sfGFP) was systematically lower than the log_2_(mKate2) for both truncated variants (Fig 4B and Table S6B). The average [log_2_(sfGFP) - log_2_(mKate2)] differences ± standard errors were 0.06 ± 0.29, - 0.34 ± 0.08, and -1.7 ± 0.4 for fCidA-fCidB, fCidB-TrA, and fCidB-TrB, respectively. The differences were significantly more pronounced for fCidB-TrB (Kruskal-Wallis rank sum test, *χ^2^* = 7.2, *df* = 2, *p*-value = 0.027), while fCidB-TrA was not statistically different from fCidA-fCidB (Wilcoxon rank sum test, W=9, *p*-value = 0.1). Note, however, that the relatively small number of replicates (due to the technical complexities of such experiments) results in low discrimination power. However, in all *f*CidB-TrA replicates, the log_2_(sfGFP) was always lower than the log_2_(mKate2). Overall, these data suggest that truncations reduce the stability of the CidB variants.

#### Full-length *cidB* correlates with CI patterns, while short *cidB* does not

*In vivo*, the time between sperm maturation and actual fertilization can take several days. Toxin stability is of paramount importance for CI: toxins must be sufficiently stable to be loaded in mature sperm and persist until the sperm-to-paternal pronucleus transition (Horard et al. 2022). Our *in cellulo* transfection analyses indicated that the long CidB variants are more toxic and more stable than the short CidB variants, thus throwing into question their respective role *in vivo*: does long CidB dictate CI patterns alone, or do short CidB variants also play a role? To answer this question, we sought correlations between male *cidB* repertoires (truncated and full-length *cidB*) and CI crossing patterns. To this end, we used our previously developed four-reference framework: males from focal lines were crossed with females from four reference lines (Atyame, Duron, et al. 2011). We focused on males infected with *w*Pip-IV, since (i) they have been extensively studied and previously crossed, showing CI pattern polymorphism (Atyame et al. 2014; Atyame et al. 2015; Bonneau, Atyame, et al. 2018; Bonneau, Landmann, et al. 2018; Bonneau et al. 2019) and (ii) they have a low *cidB* variant diversity, with only seven variants described to date (Table S3), thus making putative correlations easier to detect.

When crossed with females from the four-reference isofemale lines, males infected with *w*Pip-IV from nine different isofemale lines disclosed three distinct patterns, named crossing phenotypes 1, 2, and 3 (Table 1, (Atyame et al. 2014)). Sequencing results showed that *w*Pip in these males with three distinct crossing phenotypes harbor distinct full-length *cidB*. Furthermore, males infected with a *w*Pip of the same full-length *cidB* version have the same crossing phenotype, regardless of their short *cidB* repertoires (Table 1 and Table S3). For instance, males from lines Tab-2 and Dou-1, with the same crossing phenotype 2, strongly differ in terms of the short *cidB* repertoire, although they have an identical full-length *cidB*. Similar reasoning can be applied to all four long *cidB* identified: the same full-length *cidB* always results in the same crossing phenotype.

**Table 1.** Crosses between *w*Pip-IV infected males and females from the four reference strains. C stands for compatible and CI for incompatible; the number in brackets corresponds to the number of egg rafts examined. Three distinct crossing phenotypes are observed (1, 2, and 3) that match the long *cidB* but not the short (truncated) *cidBs* present in the repertoires.

| Males | Females |  |  |  | Crossing Phenotype | Long *cidB* | Short *cidB* repertoire |  |  |  |  |  |  |
| --- | --- | --- | --- | --- | --- | --- | --- | --- | --- | --- | --- | --- | --- |
|  | Lavar | Maclo | Slab | Ist |  |  | *cidB-IV-a1* | *cidB-IV-a2* | *cidB-IV-a3* | *cidB-IV-b1* | *cidB-IV-b2* | *cidB-IV-b3* | *cidB-IV-h2* |
| Bou-1 | C (17) | C (18) | CI (25) | C (15) | 1 | *cidB-IV-a1* | x |  |  |  |  | x |  |
| Guel-1 | C (21) | C (23) | CI (20) | C (20) | 1 | *cidB-IV-a1* | x |  |  |  |  | x |  |
| Guel-2 | C (21) | C (22) | CI (24) | C (24) | 1 | *cidB-IV-a1* | x |  |  | x |  | x |  |
| Ha | C (5) | C (10) | CI (12) | C (15) | 1 | *cidB-IV-a1* | x |  |  | x |  | x |  |
| Tab-2 | C (24) | C (18) | C (14) | C (13) | 2 | *cidB-IV-b1* |  |  | x | x |  |  | x |
| Dou-1 | C (13) | C (20) | C (13) | C (23) | 2 | *cidB-IV-b1* |  |  |  | x |  | x |  |
| Is | CI (26) | CI (53) | C (33) | C (26) | 3 | *cidB-IV-b2* | x |  |  | x | x |  |  |
| Souk-2 | CI (16) | CI (16) | C (23) | C (13) | 3 | *cidB-IV-a2* | x | x | x |  |  |  |  |
| CAA | CI (17) | CI (26) | C (30) | C (12) | 3 | *cidB-IV-a2* |  | x |  |  |  |  |  |

### Polymorphism outside the binding interface regions influences CI patterns

#### New polymorphism detected outside the interface regions

When sequencing the full-length *cidB* gene in all 73 individuals, we found new polymorphism in a 3’ region located between the binding interface regions and the DUB domain, previously described as monomorphic (Bonneau, Atyame, et al. 2018). A total of nine distinct nucleotide sequences resulting in five different amino acid sequences were identified (Fig S5). There were two groups of sequences: (i) those identical to the sequence found in *w*Pip-Pel, *cidB-Pel*, or very similar (differing by one amino acid at most), which we named *cidB-Pel-alt1* to *cidB*-*Pel-alt4*; and (ii) those identical to the sequence found in the *w*Pip strain of Buckeye (Beckmann and Fallon 2013), *cidB-Buck*, or very similar (differing by a maximum of three amino acids), which we named *cidB-Buck-alt1* to *cidB*-*Buck-alt3*. The CidB-Pel and CidB-Buck regions differ by 16 amino acids, located between amino acids 693 and 765 (nucleotides 2064 to 2352 in *cidB*, within the second pseudo PD-(D/E)XK domain), downstream of the previously described binding interface regions and upstream of the DUB domain (Fig S5). *Pel*-like *cidB* were found in 30 out of 41 lines, and *Buck*-like *cidB* in the remaining 11 lines. Both *Pel-*like *cidB* and *Buck-*like *cidB* were found in the four individuals from the Maurin and Ganges natural populations (Table S3).

We transfected S2R+ cells with a CidB variant displaying a Buck region (CidB-IV-a2-Buck) instead of the previously tested CidB-IV-a2-Pel variant. CidB-IV-a2-Buck was localized in the nucleus, and no mitosis was observed using live confocal microscopy, showing that this variant is also toxic (Fig S6).

#### *Buck* versus *Pel* 3’ polymorphism of *cidB* influences CI patterns

Previous studies showed that in males infected with *w*Pip-IV *Wolbachia*, a *cidB* downstream region named *cidB-IV-2* correlated with the ability to induce CI when crossed with Tunis *w*Pip-I females (Bonneau, Atyame, et al. 2018; Bonneau et al. 2019). The presence of this region significantly matched the CI patterns: out of 245 *w*Pip-IV isofemale lines screened, males from 77 lines were incompatible with Tunis females compared to 168 that were compatible. All 77 incompatible lines were infected with a *w*Pip with the *cidB-IV-2* region in its repertoire, whereas 159 of the 168 compatible lines did not have it. The nine compatible lines with the *cidB-IV-2* region were named “discordant lines” [19]. We PCR-screened 21 lines (15 incompatible and 6 discordant) with a *cidB-IV-2* region. *cidB-IV-2* was the full-length variant for all the tested strains, thus ruling out the role played by truncation in the discordant lines (Table S1). We then acquired their long *cidB* sequence using direct Sanger sequencing and found that all the discordant lines had a Buck-like 3’ polymorphic region, whereas the others had the Pel-like one (Table 2).

**Table 2.** Crossing phenotypes between *w*Pip-IV infected males and females from the Tunis isofemale line (*w*Pip-I) correlate with the polymorphism in the 3’ region of *cidB*. All strains with the *cidB-IV-2* region have a complete *cidB-IV-2* version. Sanger sequencing of the long *cidB* revealed polymorphism in its 3’ region similar to either the Pel or Buck regions. The Buck region is present in all discordant lines and absent (marked as an A) from all incompatible lines.

| Type | Males | Tunis | Long *cidB-IV-2* | *Buck-like* in long *cidB* |
| --- | --- | --- | --- | --- |
| Discordant | Hamra11 | C | x | x |
|  | Ich12 | C | x | x |
|  | Ich30 | C | x | x |
|  | Michele26 | C | x | x |
|  | Ut44 | C | x | x |
|  | Ut63 | C | x | x |
| Incompatible | Hamra17 | IC | x | A |
|  | Hamra21 | IC | x | A |
|  | Ich03 | IC | x | A |
|  | Ich09 | IC | x | A |
|  | Ich21 | IC | x | A |
|  | Ich24 | IC | x | A |
|  | Ich28 | IC | x | A |
|  | Istanbul | IC | x | A |
|  | Ut50 | IC | x | A |
|  | Souk2 | IC | x | A |

The influence of this 3’ region on CI crossing types is further supported by other crossing results involving males infected with *w*Pip-II. Males from three lines (Lavar, Australie, and LaCartara1) with the exact same short *cidB* repertoire differed in terms of the induced CI patterns (Table 3). We found that their full-length *cidB* was identical on the binding interface regions but differed in the 3’ region (Australie and LaCartara1 had a Buck-like *cidB* and Lavar a Pel-like *cidB*).

**Table 3.** Polymorphism in 3’ of *cidB* perfectly matches differences in crossing patterns. Males infected with *w*Pip-II were crossed with females from the four reference strains, showing two distinct *mod* cytotypes (Lavar vs. the other strains). Lavar has a Pel-like *cidB*, whereas the two other strains have a Buck *cidB*.

|  | Males | Females |  |  |  | Phenotype | Long *cidB* | Short *cidB* |  | 3' region |
| --- | --- | --- | --- | --- | --- | --- | --- | --- | --- | --- |
|  |  | Lavar | Maclo | Slab | Ist |  |  | *cidB-II-u1* | *cidB-II-u3* |  |
| **wPip-II** | Lavar | C (8) | C (36) | IC (30) | IC (40) | 1 | *cidB-II-u1* | x | x | Pel |
|  | Australie | C (20) | C (9) | IC (34) | C (11) | 2 | *cidB-II-u1* | x | x | Buck |
|  | Cartara1 | C (38) | C (23) | IC (42) | C (18) | 2 | *cidB-II-u1* | x | x | Buck |

## Discussion

*Wolbachia* induce highly variable CI crossing patterns in *Culex pipiens* (Duron et al. 2006; Atyame et al. 2014), previously linked to the polymorphism of *cid* genes in the CidA-CidB interaction regions (Bonneau et al. 2019; Sicard et al. 2021; Namias et al. 2024). More specifically, the presence or absence of *cidA* variants (antidotes) correlates with distinct *rescue* patterns in females, while the presence or absence of *cidB* variants (toxins) correlates with distinct *modifications* in males. Although correlations between *cid* repertoires and crossing patterns were previously found, the relation between *cid* repertoires and crossing patterns has not yet been fully deciphered.

Here, we uncovered a further layer of polymorphism in *cidB*, which consequently improved our understanding of CI patterns. By studying numerous *cid* repertoires, we revealed their architecture and evolutionary origin for the first time.

### High polymorphism in the CidA-CidB binding interface regions results from recombination

By sequencing more *w*Pip lines, we described even more polymorphism and confirmed the high variability of *cid* genes in CidA-CidB interface regions. *cid* repertoire acquisitions from natural populations showed that *cid* amplification and diversification are not a laboratory oddity. We demonstrated that *cid* polymorphism mostly results from recombination. *cid* variants are composed of numerous recombination blocks for which we found two alternative alleles (Fig 1 and Fig S1). A plausible explanation is an ancestral state with two distinct *cidA-cidB* pairs within a single genome followed by numerous recombination steps, resulting in many recombination blocks as well as the highly complex variants that we observe today.

*w*Pip-infected *Culex* mosquitoes exhibit the most complex CI crossing types described to date (Atyame et al. 2014). These patterns were found to correlate with the existence of five phylogenetic groups within *w*Pip, with mosquitoes infected with *Wolbachia* from the same phylogenetic group being largely compatible (Atyame et al. 2014). Surprisingly, most *cid* recombination blocks observed were common to all *w*Pip phylogenetic groups. Only three block alleles were found to be group-specific (one *cidA* block and two *cidB* blocks), while all of them were specific to *w*Pip-IV, a group previously shown to be strongly incompatible with other *w*Pip groups (Atyame et al. 2014; Bonneau, Atyame, et al. 2018; Bonneau et al. 2019; Sicard et al. 2021).

Although block alleles are shared among *w*Pip groups, variants (i.e., allelic associations between blocks) are specific to one group, with a few rare exceptions. These results suggest that some block allele associations, or even whole variants, are responsible for compatibility as opposed to blocks alone.

### Truncations in *cidB* shape CI patterns

#### A single full-length *cidB* is key to CI crossing types

Numerous *cidA* and *cidB* variants are found in each *w*Pip genome, thus making it difficult to decipher the toxin/antidote (TA) interactions underlying the different CI patterns. Here, we uncovered a further layer of polymorphism by showing that within each *w*Pip genome, all but one *cidB* were truncated, missing their 3’ end, including the DUB domain previously shown to be key for CI (Beckmann et al. 2017; Horard et al. 2022). A correlative approach between variants and CI patterns within the phylogenetic group *w*Pip-IV strongly suggests that only the single full-length *cidB* variant plays a role in CI crossing patterns (Table 1). This simplifies the conception of the TA mechanism in *Culex*: a single toxin, and not multiple ones, would have to be rescued to make a cross compatible (Fig. 5).

**Figure 5.**
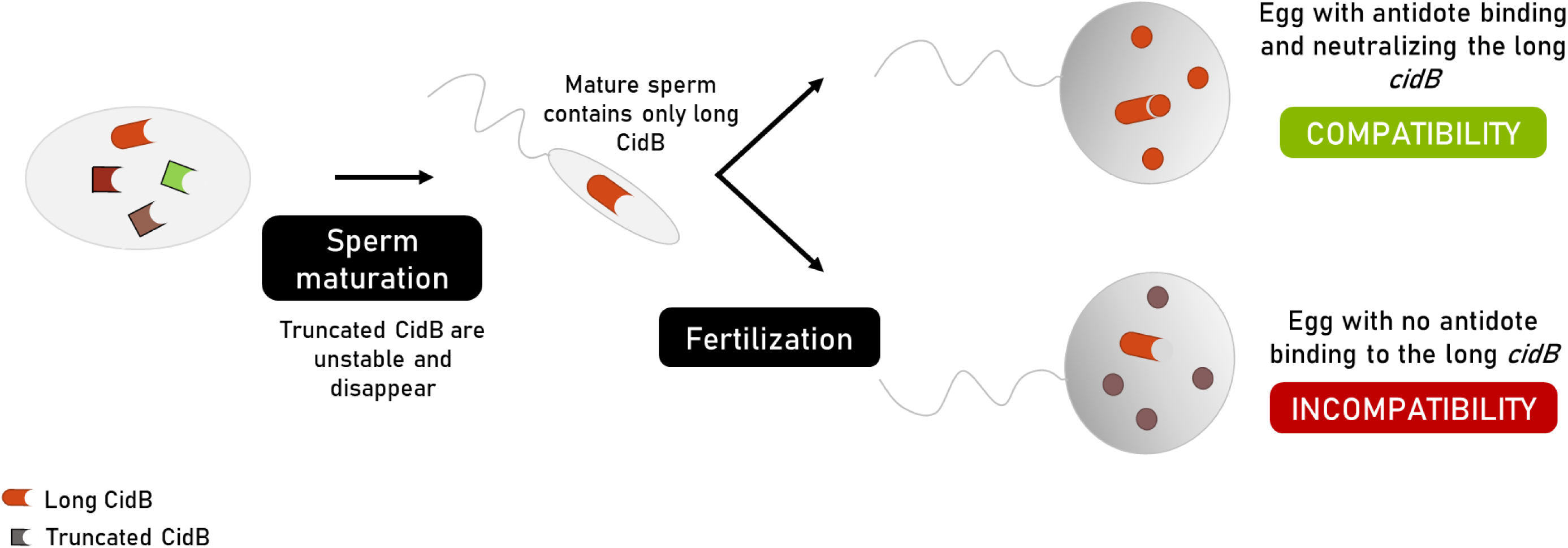
Only long *cidB* plays a role in CI. *Wolbachia* is eliminated over the course of sperm maturation. One hypothetical explanation for the role played by the sole long CidB in CI is that truncated CidB, which are unstable, disappear as sperm matures. In mature sperm, the only CidB remaining is thus the long CidB.

The first hypothesis to explain this result is that the truncated variants are not expressed. This hypothesis can, however, be ruled out, as previous work (before the discovery of truncations) showed that all the variants present in the *cidB* repertoire of a given strain occurred in the corresponding cDNA (Bonneau, Atyame, et al. 2018). By expressing truncated variants in *Drosophila* cells, we showed that (i) they were toxic, even though toxicity is much lower than that of full-length CidB; (ii) this toxicity could be rescued by the same *cidA* as with the full-length *cidB*, showing that truncation did not influence the interaction zones; and (iii) they exhibited potentially reduced stability compared to the full-length *cidB*s, likely resulting from the truncation of their DUB domain, a domain known to be associated with stabilizing properties (Clague et al. 2012; Harumoto 2023; Terretaz et al. 2023).

Previous cytological analyses of paternal *Wolbachia* transmission suggest an explanation as to why only complete variants influence CI patterns: *in vivo*, *Wolbachia* are removed from maturing sperm cells (Bressac and Rousset 1993), and thus only CidB proteins persist and are transmitted to the egg with sperm (Horard et al. 2022); if truncated proteins are unstable, they may be degraded before fertilization can occur, so that only full-length CidB can be transmitted to the egg and induce CI phenotypic effects (Fig 5).

#### Origin of truncations: Horizontal transfers and recombination

Truncated *cidB* are organized into palindromes, which are absent from the reference *w*Pip-Pel genome draft assembly but are present in *w*Pip-JHB contigs. These palindromic structures could have been acquired in the *w*Pip genomes through (i) vertical transmission from a common ancestor or (ii) horizontal transfers from another *Wolbachia* – *w*Naev; the *Wolbachia* infecting the moth *Rhopobota naevana* is the best candidate based on currently available data. Such horizontal transfers have frequently been described in *Wolbachia* genomes (e.g., (Martinez et al. 2021; Tan et al. 2024)). A few hypotheses have been put forward in the literature (e.g., transfer of *Wolbachia* through predation or a shared nutritional source (Le Clec’h et al. 2013; Ahmed et al. 2016; Li et al. 2017), transfer through phages (Kaur et al. 2022) or insertion sequences (Cooper et al. 2019)), although experimental evidence is still scarce.

The lack of homologous palindromic structures in *Wolbachia* strains closely related to *w*Pip and the presence of an insert matching the *cidB* of a distantly related strain, wNaev, strongly suggest that palindromes are not the ancestral *cid* architecture in *w*Pip. *w*Pip palindromes more likely arose from a horizontal transfer followed by genomic rearrangements (by some unidentified mechanism) of *cid* pairs after transfer. It would appear that the truncated parts of both *cidB* involved in the palindrome P1 are located next to each other in a different *w*Pip genomic region, suggesting that the truncation of palindromic *cidB*s was caused by a single event concomitant with the creation of the palindrome. Palindrome P2, which is not fixed in *w*Pip, could result from the rearrangement of palindrome P1 and replacement of *cidB-TrB* by *cidB-TrC*. Within-palindrome rearrangements must have occurred, as all tested strains share palindrome P1 without necessarily sharing *cidB* variants.

Among the 138 *Wolbachia* genomes analyzed here, a single palindrome was identified in only one genome, namely *w*Naev, from which a *cidB* was likely transferred. The high quality of the genome sequences used here makes assembly problems unlikely. It could be envisaged that such palindromic structures favor gene rearrangements. Indeed, some palindromic structures (also known as inverted repeats) have already been shown to play a role in recombination (e.g., with small palindromes forming hairpins in bacteria (Bikard et al. 2010)) or in genetic instability and gene amplifications in a more general way (e.g., in humans (Tanaka et al. 2006) and in bacteria (Achaz et al. 2003)).

#### Maintenance of truncated *cidB*

Crossing data suggest that truncated *cidB* genes play no role in CI crossing phenotypes that correlate perfectly with the identity of the full-length *cidB*, regardless of the short *cidB*. At least three alternative scenarios may explain the maintenance of these truncated variants in all sequenced *w*Pip repertoires: (i) these variants are neutral or under negative selection, but pseudogenization is too recent for genes to have been lost; (ii) truncated *cidB* genes are under positive selection; or (iii) truncated *cidB* genes are neutral *per se* but kept by hitchhiking. The first hypothesis is plausible, as *w*Pip divergence is recent, with *w*Pip genomes showing low diversification, except for highly repeated regions such as *cid* genes or genes used for the specific *w*Pip MLST (Atyame, Delsuc, et al. 2011). It is possible that pseudogenes did not have time to be eliminated. We can also imagine that truncated *cidB* play a role in a non-CI process such as regulating the density of *Wolbachia* through autophagy interactions (Deehan et al. 2021). Alternatively, truncated *cidB* may have no advantage *per se* but only be kept by hitchhiking. Modeling studies previously showed that only *cidA* genes were under selection in randomly mating populations (Turelli 1994). Maintaining a high diversity of *cidA* genes could be advantageous at the population level, because losing one *cidA* could result in a loss of compatibility with surrounding lines (as we recently showed in (Namias et al. 2024)). Maintaining *cidA* could also be advantageous at an individual level: the cell experiment suggests that truncated *cidB* can still be toxic at the cell level, meaning that *cidA-*truncated *cidB* pairs could still act as an addictive module within an individual host similar to conventional toxin-antitoxin systems that are not involved in CI. In both cases, maintaining the *cidA* repertoire is under positive selection, and *cidB* could be maintained due to their tight linkage with the *cidA* genes.

#### Polymorphism outside the CidA/CidB binding interaction regions influences CI patterns

In addition to truncations, we identified polymorphism in the 3’ region of the *cidB* gene, which corresponds to the second pseudo PD-(D/E)X/K domain. This polymorphism was previously missed due to the sequencing method, which only amplified a restricted part of the *cidB* gene (Bonneau, Atyame, et al. 2018). This polymorphism, located outside the previously described CidA-CidB binding interface regions, influences CI crossing phenotypes and notably solves the discrepancies in CI patterns, which could not be explained by *cid* polymorphism in the binding regions alone. Two main groups of sequences were described in this 3’ region: *Pel-*like, previously described in the reference *w*Pip-Pel genome, and *Buck-*like, similar to the allele present in *w*Pip-Buckeye (Beckmann and Fallon 2013). Experiments in *Drosophila* cells showed that both full-length CidB-Pel and Buck behave in a similar way: they are localized in the nucleus and are toxic. CidB-Buck are also able to induce CI *in vivo*, as strains shown here to have a long *cidB-Buck* were previously demonstrated to induce CI in crosses (Atyame et al. 2014).

This 3’ region was shown to be required for CI onset: its deletion prevents CidB toxicity and nuclear import in *Drosophila* cells (Terretaz et al. 2023). Because it affects neither the interface region nor the DUB region required for Cid upload in the sperm, the influence of Buck/Pel polymorphism questions the toxin-antidote framework, which defines toxicity or rescue by the sole binding (or not) of the toxin and the antidote.

We have two alternative hypotheses to explain the role of this 3’ region in CI pattern polymorphism: (i) this region influences protein folding and changes the interaction regions, or (ii) CidB interacts differently with downstream effectors depending on the identity of this specific 3’ sequence. A recent study showed CidB homologs in *w*Mel (infecting *Drosophila melanogaster*) are nucleases whose function relies on the presence of a QxxxY motif (Kaur et al. 2024). The difference between CidB-Buck and CidB-Pel could be explained by a change in this nuclease region, but no QxxxY motif was observed in CidB-Pel or CidB-Buck. However, another motif not yet identified may be responsible. New experiments to elucidate the role of this region in CidA-CidB interactions should be explored, such as the strength of binding between different toxins and antidotes.

Our previous acquisitions of *cid* repertoires uncovered a huge diversity of *cid* variants. Given this high complexity, it was impossible to link the identity of *cid* genes with CI patterns. Here, using other sequencing methods, we showed that all but one *cidB* gene are truncated and that the long *cidB* seems to be the only one involved in CI patterns, which simplifies the understanding of the complex CI patterns in *Culex pipiens*. We show that truncations likely reflect the horizontal transfers of *cid* genes among *Wolbachia*, thus further fueling the high rate of horizontal transfers in *Wolbachia* genomes.

## Methods

### Mosquitoes used in the study

Table S1 provides a list of all the lines used in this study along with their respective geographic origins. Unless mentioned otherwise, all mosquitoes used here come from isofemale lines, i.e., the progeny of a single egg raft and thus from a single female. Mosquitoes were reared in 65 dm^3^ screened cages in a single room maintained at 26°C under a 12h light/12h dark cycle. Larvae were fed with a mixture of shrimp powder and rabbit pellets, and adults were fed on honey solution. Females were fed with turkey blood using a Hemotek membrane feeding system (Discovery Workshops, UK) to enable them to lay eggs.

### Crosses

For each cross performed, around 50 virgin females were put in a cage with around 50 virgin males. After five days spent in the cage, females were fed a blood meal. Shortly after laying, the egg rafts were moved to 24-well plates. Cross compatibility was assessed 2 days after egg laying: the cross was classified as compatible if eggs had hatched and as incompatible if none had hatched. Egg rafts with a null hatching rate were mounted on a slide and checked with a binocular magnifier to ensure that they were fertilized and that the null HR resulted from CI as opposed to the absence of fertilization.

### Repertoire analysis

#### Nanopore sequencing of PCR products

All repertoires were sequenced following (Namias et al. 2023). Briefly, DNA was extracted following the CTAB protocol (Rogers and Bendich 1994). *cid* genes were amplified with a specific PCR: a single PCR was used for *cidA*, encompassing all the variable regions (Bonneau, Atyame, et al. 2018), whereas two PCRs were used for *cidB*, one PCR amplifying the variable regions (“short” *cidB*) and another amplifying the complete *cidB* variant (“long” *cidB*). PCR products were then purified. In each well, *cidA* and *cidB* PCR products were pooled in an equimolar mix. All PCR products were sequenced using MinION technology by the Montpellier GenomiX platform. All primers and PCR conditions are found in Table S4.

#### Direct sequencing of long *cidB*

Direct Sanger sequencing was also used for long *cidB* analyses. This was performed (i) to check the existence of a single long *cidB* in each individual and (ii) to confirm which allele of the long *cidB* (Pel/Buck) was present in lines from *w*Pip-IV bearing *cidB-IV-2*. To do so, a fragment was amplified by PCR using (i) the “long *cidB*” primers or (ii) the “long *cidB-*IV-2’ primers (Table S4). These fragments were then Sanger sequenced.

#### Sequencing and analyses of Nanopore long reads

High molecular weight genomic DNA was isolated from 10 females for each line using the Qiagen Genomic-tip 20G kit following the manufacturer’s protocol for insects. DNA libraries were prepared using the Ligation Sequencing Kit (SQK-LSK109) and Native Barcoding Expansion Kit (EXP-NBD104) and then sequenced on a Minion Mk1B using a FLO-MIN106D flow cell with R9.4.1 chemistry. Nanopore reads were basecalled with GuPPy v3.3.0 (Sherathiya et al. 2021) and assembled with Canu v2.2 (Koren et al. 2017). The assembly was corrected with Illumina reads via Pilon v1.24 (Walker et al. 2014). Putative *cid* genes were identified by blasting *cid* homologs against the assembly. The contigs containing truncated *cidB* were short and could not be extended. We continued our investigations and found that the truncated *cid* was in the center of a long palindromic sequence in which no Nanopore reads covered both the *cid* and unique sequence beyond the palindromic region, which explains why the contig could not be extended.

#### Specific PCRs

Several PCRs were used to test for the presence or absence of a specific target region. In all cases, primers were designed using a positive and a negative control and following (Namias et al. 2023) (Fig S3) to ensure that none of the other variants present could be amplified. All primers, along with the PCR conditions, are outlined in Table S4.

#### Analysis of the alternative *cidB* allele in “discordant” strains

The presence of the *cidB-IV-2* downstream region was checked using a specific PCR (“Short *cidB-IV-2*” in Table S4, (Bonneau et al. 2019)). Then, we tested whether the complete *cidB* contained this region using a PCR anchored in a specific *cidB-IV-2* region on one side and in the 3’ region of *cidB* on the other side (absent from truncated variants, named “Long *cidB-IV-2*” in Table S4). PCR products were purified using the Agencourt Ampure PCR purification kit (Agencourt) and directly sequenced with an ABI Prism 3130 sequencer using the BigDye Terminator Kit (Applied Biosystems).

#### Recombination analysis

The existing recombination blocks were identified using RDP4 (Martin et al. 2015). We confirmed the existence of a recombination block when it was validated by at least four methods.

#### Phylogenetic analysis and palindrome distribution in *Wolbachia* genomes

A dataset of 138 publicly available genomes belonging to the *Wolbachia* supergroups A and B was used to build a phylogeny of the symbiont. Overall, 80% (109/138) of the genomes came from the “Darwin Tree of Life” project (https://www.sanger.ac.uk/programme/tree-of-life/) and were assembled using high-quality PacBio Hifi long-read technology, which facilitated the assembly of prophage and other repetitive regions (including *cid* genes) commonly found in *Wolbachia* genomes (Vancaester and Blaxter 2023). Roary v3.11.2 (Page et al. 2015) was used to identify the sequence of 292 single-copy genes with a minimum of 95% identity and shared by >99% of genomes and to generate a concatenated gene alignment. The nucleotide alignment was then used to build a *Wolbachia* phylogeny using RaxML-NG v.1.0.2 (Kozlov et al. 2019) with the GTR+G substitution model and 100 bootstrap replicates.

The presence of *cif* palindromes in the corresponding *Wolbachia* genomes was analyzed using TBLASTN and a set of representative *cifA* and *cifB* protein sequences from type I to V as queries. Positive hits were visually inspected along the *Wolbachia* genomes using the Artemis genome browser v.16.0.0 (Carver et al. 2012). All the detected *cif* genes and pseudogenes were manually annotated within Artemis, and their amino acid sequence was extracted. Sequences were then aligned with the PROMALS3D web server (http://prodata.swmed.edu/promals3d/), and weakly conserved regions were filtered out from the alignment using trimAl v1.5.0 with the automated 1 setting (Capella-Gutiérrez et al. 2009). The curated alignments of *cifA* and *cifB* homologs were used to build phylogenies with RAxML-NG in order to determine the *cif* type of each homolog (I to V).

### Tests of toxicity and stability in cells

#### Experiments

All assays were performed following (Terretaz et al. 2023). Briefly, *D. melanogaster* S2R+ cell lines obtained from the Drosophila Genomics Resource Center were cultured in Schneider’s *Drosophila* medium (Dutscher #L0207-500) supplemented with 10% Fetal Bovine Serum (Dutscher #S1810-500) at 25°C.

A synthetic cassette (Genscript) containing the mkate2-T2A-sfGFP block was inserted in the multiple cloning site of a *Drosophila* cell vector based on the pMT-V5-HisC (Invitrogen #V412020), which was modified to have an Actin5C promoter. *cidA-IV-delta(1)-1* and *cidB-IV-a-2* genes were synthesized after codon optimization for expression in *D. melanogaster* cells (Genscript). The third 73 bp intron of the *D. melanogaster* nanos (nos) gene was inserted close to the 5’ end of CidB to avoid toxic leak encountered in E. coli (Horard et al. 2022). All plasmids were obtained by Gibson cloning using the NEBuilder Hifi DNA Assembly kit (NEB #E5520S) and verified by Sanger sequencing.

For live microscopy, cells were plated in 35 mm glass bottom dishes (Cellvis #D35-20-1.5-N) and transfected with Lipofectamine 3000 (Invitrogen #L3000008) with 500 ng of purified plasmid DNA according to the manufacturer’s instructions. Then, 24h post-transfection, 48h time-lapse recordings were performed on transfected cells to evaluate the toxicity of Cid variants. Percentages of mitotic and apoptotic events of transfected cells were then compiled from two independent transfection experiments. For each variant, at least 1,000 transfected cells were counted. In addition, transfected cells were observed by confocal microscopy between 24 and 48h after transfection to determine the localization of Cid variants.

Cell stability and viability assays were conducted as previously described using flow cytometry (Horard et al. 2022). Briefly, cells were analyzed in three biological replicates 2 and 4 days after transfection, and a growth rate was calculated according to the following formula: Log2 (*x* at day 4 / *x* at day 2), where *x* is the proportion of fluorescent cells at the given time-point. A fold change equal or superior to 0 was observed when transfected cells grew at a similar rate to non-transfected cells. By contrast, a negative fold change reflects slower growth or cell death between day 2 and day 4. The instability of CidB mutants was deduced from the discrepancy between a decreasing sfGFP fluorescence level and a stable or increasing free mKate2 fluorescence level. Data were acquired using a Novocyte ACEA cytometer and analyzed with the NovoExpress (ACEA) software.

### Statistical analysis

All computations were performed using R 4.4.0 (Team 2013).

Variability in the proportion of mitosis versus apoptosis was analyzed using a generalized linear mixed model (GLMM), with the number of cells in mitosis over the number of cells in mitosis or apoptosis as a dependent variable, and the construct (four levels: fCidA-fCidB, fCidB, fCidB-TrA, or fCidB-TrB) as the independent variable. Mixed effects were used to account for differences among replicates, while the error parameter followed a binomial distribution. We used the package lme4 (Anon 2024). Computed models were simplified by testing the significance of the different terms using likelihood ratio tests (LRT), starting from the higher-order terms. Factor levels of qualitative variables, whose estimates (using LRTs) were no different, were grouped as described by Crawley (Crawley 2007).

FACS data cannot be immediately compared from one replicate to another, as GFP or mkate2 counts strongly depend on the cell state, which varies among replicates and experiments. We directly analyzed the Δ_GFP-mKate2_ = log_2_(sfGFP) -log_2_(mKate2) difference between constructs. As their distribution is unknown, we used a non-parametric approach with a Kruskal-Wallis rank sum test, with the Δ_GFP-mKate2_ as the dependent variable, and the construct (three levels: fCidA-fCidB, fCidB-TrA or fCidB-TrB) as the independent variable. When a significant effect was found, we used Wilcoxon rank sum tests between the closest constructs to identify the significant differences.

### Protein structure

Protein domains of the *cidB* from *w*Pip-Naev were predicted using the HHPred webserver (Söding et al. 2005) following (Lindsey et al. 2018), with default parameters and the following databases: SCOPe70 (v.2.08), Pfam (v.37), SMART (v6.0), and COG/KOG (v1.0).

## Acknowledgments

We thank Nicole Pasteur and Michael Turelli for their comments and discussions on the manuscript. We thank Brandon S. Cooper for the Nanopore sequencing of the Tunis and Harash lines and for supporting William Conner through a grant from the US National Institutes of Health (R35GM124701). We also thank Tim Wheeler for contributing to this work. We thank Infravec for preliminary sequencing data and the GenSeq platform for Qubit and direct Sanger sequencing. The Nanopore sequencing data used here were generated on the Montpellier GenomiX platform. This project was funded by the MUSE project (reference ANR-16-IDEX-0006).

